# Interaction between variants in *CLU* and *MS4A4E* modulates Alzheimer’s disease risk

**DOI:** 10.1101/028951

**Authors:** Mark T. W. Ebbert, Kevin L. Boehme, Mark E. Wadsworth, Lyndsay A. Staley, for the Alzheimer’s Disease Neuroimaging Initiative, Alzheimer’s Disease Genetics Consortium, Shubhabrata Mukherjee, Paul K. Crane, Perry G. Ridge, John S. K. Kauwe

## Abstract

**INTRODUCTION**: Ebbert et al. reported gene-gene interactions between rs11136000-rs670139 *(CLU-MS4A4E)* and rs3865444-rs670139 *(CD33-MS4A4E).* We evaluate these interactions in the largest dataset for an epistasis study.

**METHODS**: We tested interactions using 3837 cases and 4145 controls from ADGC using meta-and permutation analyses. We repeated meta-analyses stratified by *APOEε4* status, estimated combined OR and population attributable fraction (cPAF), and explored causal variants.

**RESULTS**: Results support the *CLU-MS4A4E* interaction and a dominant effect. An association between *CLU-MS4A4E* and *APOEε4* negative status exists. The estimated synergy factor, OR, and cPAF for rs11136000-rs670139 are 2.23, 2.45 and 8.0, respectively. We identified potential causal variants.

**DISCUSSION**: We replicated the *CLU-MS4A4E* interaction in a large case-control series, with *APOEε4* and possible dominant effect. The *CLU-MS4A4E* OR is higher than any Alzheimer’s disease locus except *APOEε4, APP,* and *TREM2.* We estimated an 8% decrease in Alzheimer’s disease incidence without *CLU-MS4A4E* risk alleles and identified potential causal variants.

## 1. Introduction

Alzheimer’s disease (AD) is a complex neurodegenerative disease, and is the third leading cause of death in the United States [1]. AD is characterized by the accumulation of amyloid plaques and neurofibrillary tangles in the brain. Many genetic loci exist that modify AD risk, but collectively, they explain only a fraction of AD’s heritability [2] and are not diagnostically useful [3,4]. Rare variants with large effects and epistatic interactions may account for much of the unexplained AD heritability, but are largely unknown due to limitations in traditional GWAS studies. While rare variant and epistatic effects on AD are poorly understood, recent studies suggest that gene-gene interactions play a critical role in AD etiology and progression [3,5–7].

A previous study [3] reported evidence of two gene-gene interactions that increase AD risk. Specifically, Ebbert et al. reported interactions between rs11136000 C/C (*CLU*; minor allele = T, MAF = 0.38) and rs670139 G/G (*MS4A4E;* minor allele = T, MAF = 0.38) genotypes (synergy factor (SF) = 3.81; p = .016), and the rs3865444 C/C (*CD33;* minor allele = A, MAF = 0.21) and rs670139 G/G *(MS4A4E)* genotypes (SF = 5.31; p = .003). All three variants have been implicated in numerous AD GWAS studies [8–13] and are on the “AlzGene Top Results” list [14], which summarizes the most established genes associated with AD.

*MS4A4E* and *CLU* were recently replicated in a large meta-analysis of 74046 individuals, but *CD33* did not replicate [15]. Despite *CD33* failing to replicate, several studies demonstrated that *CD33* is involved in AD-related pathways and pathology, giving convincing evidence that *CD33* is somehow involved in AD. Three specific studies demonstrated that *CD33* alters monocyte function, amyloid uptake, and that *CD33* expression is associated with clinical dementia ratings [16–18]. rs3865444 is located in the 5’UTR of *CD33.*

The association between *CLU* and AD status has been strongly established by both genetic and biological data. Recent studies demonstrated that rs11136000—an intronic SNP within *CLU*—is associated with AD-related pathology in healthy individuals including neural inefficiency [19] and decreased white matter integrity [20].

*MS4A4E* is a member of the membrane-spanning 4-domains subfamily A, but little else is known about the gene. However, rs670139—located in the *MS4A4E* 3’UTR according to gene model XM_011545416.1—is consistently associated with AD [15,18,21].

In this study, we attempted to replicate these gene-gene interactions using the largest dataset used in an epistasis study, to date [22]. We performed an independent meta-analysis of datasets from the Alzheimer’s Disease Genetics Consortium (ADGC) using 3837 cases and 4145 controls, followed by a combined meta-analysis that included the original Cache County results [3] with an additional 326 cases and 2093 controls. We also tested for dosage or dominant effects and an *APOEε4* effect. Finally, we explored possible causal variants using whole-genome sequence data from the Alzheimer’s Disease Neuroimaging Initiative (ADNI).

## 2. Methods

### 2.1. Data description

We used SNP data from the ADGC, which consists of 32 studies collected over two phases and includes 16000 cases and 17000 controls. All subjects are self-reported as being of European American ancestry. More information about this dataset can be found in Naj et al. [8] and the ADGC data preparation description [23].

Genotype data from 2419 individuals from the Cache County Study on Memory Health and Aging were also used in this study. The full cohort of 5092 individuals represented approximately 90% of the Cache County population aged 65 and older when the study began in 1994 [24]. The Cache County data consists exclusively of individuals of European American ancestry. Exactly 2673 individuals were excluded from the original Cache County analysis because of incomplete genotype or clinical data [3]. Additional information on this dataset can be found in previous reports [3,24].

Whole-genome data from 747 (223 controls, 195 cases, 329 MCI) individuals were used in this article and were obtained from the ADNI database (adni.loni.usc.edu). ADNI is a large collaboration from several academic and private institutions, and subjects have been recruited from over 50 sites across the U.S. and Canada. Currently, over 1500 adults (ages 55 to 90) participate, consisting of cognitively normal older individuals, people with early or late MCI, and people with early stage AD. For up-to-date information, see www.adni-info.org.

### 2.2. SNP data preparation and statistical analysis

As gene-gene interactions are challenging to identify and replicate, we used the highest quality data possible. For each ADGC dataset, we filtered SNPs imputed with low information (info < 0.5) and converted the IMPUTE2/SNPTEST format files to PLINK format, using PLINK v1.90b2i [25,26]. We used the default PLINK uncertainty cutoff of 0.1, meaning any imputed call with uncertainty greater than 0.1 was treated as missing. We included SNPs with a missing genotype rate less than 0.05 and individuals with a missing rate less than 0.01. We then extracted the SNPs of interest: rs3865444 *(CD33),* rs670139 *(MS4A4E),* and rs11136000 *(CLU)* and tested Hardy-Weinberg equilibrium [27,28]. Using R v3.1.1 [29], we excluded samples without complete data for all covariates including age, gender, case-control status, *APOEε4* dose, and the two SNPs being tested in the corresponding interaction. Entire datasets missing the respective SNPs or covariates after data cleaning were excluded from further analysis. The requirement of complete data for both SNPs and all covariates is necessary for this analysis. Unfortunately, this requirement led to the exclusion of 23 and 24 entire datasets for the *CD33-MS4A4E* and *CLU-MS4A4E* interactions, respectively. We also excluded the ADC1 dataset because it contained only one AD case, likely making it biased.

Following data preparation, we tested the individual interactions in each dataset using logistic regression. We defined the R models as “case_control ~ rs3865444 + rs670139 + rs3865444:rs670139 + apoe4dose + age + sex” and “case_control ~ rs11136000 + rs670139 + rs11136000:rs670139 + apoe4dose + age + sex” for the *CD33-MS4A4E* and *CLU-MS4A4E* interactions, respectively. Case-control status, SNPs, and sex were coded as factors, age was numeric, and apoe4dose was an ordered factor from 0-2.

Using results from each study, we performed a meta-analysis to test replication across the ADGC datasets using METAL (version 2011-03-25) [30], and performed a second meta-analysis including the original Cache County results to provide synergy factor and odds ratio estimates from the largest number of samples possible. We tested the originally reported interactions and heterozygous interactions (rs11136000 C/C—rs670139 G/T and rs3865444 C/C—rs670139 G/T) to test for potential dosage or dominant effects based on suggestive evidence found in the original Cache County study (Supplemental Table 1). We assessed whether there is a dosage or dominant effect based both on whether the heterozygous interaction is significant and a t-test comparing two means. Specifically, we tested for a significant difference between the homozygous and heterozygous effect sizes. A significant difference would suggest a dosage effect, whereas an insignificant difference would suggest the effect might be dominant.

Following the meta-analyses, we performed a permutation analysis with 10000 permutations for interactions that replicated independently. For each ADGC dataset, we randomly permuted case-control status across all individuals, tested the interaction, and reran the meta-analysis. We stored the p-values from each of the 10000 meta-analyses and calculated the empirical p-value by finding the original p-value’s rank in the distribution of p-values divided by the number of permutations. We also calculated the combined population attributable fraction (cPAF) as previously described [3,8].

Results are represented using both odds ratios and synergy factors [6,31] and their associated 95% confidence intervals and p-values. Synergy factors represent the ratio between the *observed* and *expected* odds ratios for the two interacting SNPs (Equation 1). The *expected* odds ratio for the interaction assumes there is no synergy between the SNPs (i.e., the SNPs are independent) and equals the product of the individual odds ratios (the denominator of Equation 1) [6,31]. Essentially, the synergy factor measures how strongly the *observed* and *expected* odds ratio relationship deviates from linearity, as the synergy factor deviates from one. A synergy factor equal to one suggests no synergy; rather, there is no evidence of statistical epistasis.

**Equation 1.**
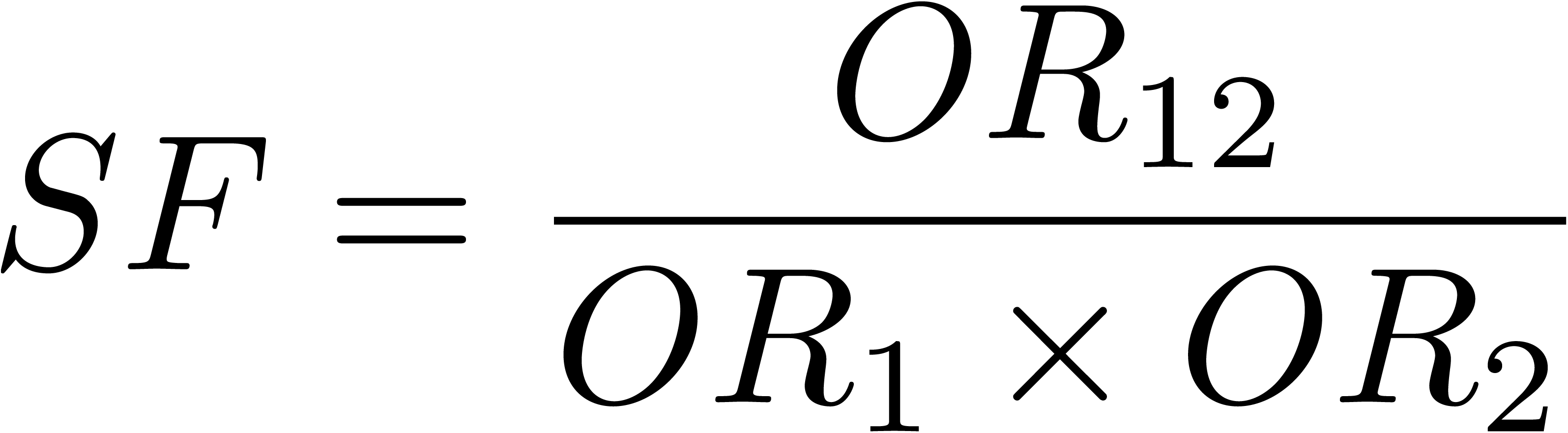
The synergy factor describes the relationship between the expected odds ratio (denominator) and the observed odds ratio (numerator) for interacting variants. The expected odds ratio (denominator) assumes that both variants are independent (i.e., there is no synergistic, or non-linear effect on the phenotype) while the observed odds ratio (numerator) is the actual effect. A synergy factor that deviates from 1 indicates a statistical interaction between the variants that affects the phenotype.

Because synergy factors less than 1 can be challenging to interpret, we present interaction synergy factors in the direction greater than 1. Consequently, we performed all interaction analyses using each gene’s homozygous minor allele as the reference group, which is opposite the direction standardly used in genome-wide association studies. This also has the added advantage that the interaction’s odds ratio is presented in the risk direction for easy comparison to top AD risk loci. To calculate the interaction’s odds ratio, we used each SNPs individual odds ratio as previously reported in a larger dataset [21], *but we had to invert the individual odds ratios to be the same direction as our analyses.* We then calculated the interaction’s *observed* odds ratio (Equation 1) using the inverted odds ratios. We also estimated each synergy factor’s 95% confidence interval using rmeta [32].

Based on results from the interaction replication, we performed a synergy factor analysis using the Cortina-Borja synergy factor Calculator [31] to test for an *APOEε4* effect for the *CLU-MS4A4E* interaction. Specifically, we stratified the combined ADGC and Cache County data by *APOEε4* status and tested for an association between the interaction and case-control status within each stratum. Alleles rs11136000 C and rs670139 G were used as the exposed groups.

### 2.3 Exploring causal variants

As a follow up analysis, we explored causal variants for replicated interactions using 747 (223 controls, 195 cases, 329 MCI) ADNI whole genomes that were sequenced, aligned to hg19, and variants identified by Illumina using their internal analysis procedure. We used linkage disequilibrium, Regulome DB (accessed November 2014) [33], and functional annotations from wAnnovar [34] to isolate SNPs of interest. We first extracted all SNPs within approximately 50 kilobases of each SNP of interest, calculated linkage disequilibrium using Haploview [35], and retained all SNPs with a D’ ≥ 0.99. Using Regulome DB and wAnnovar, we annotated each remaining SNP for: (1) known regulation and functional effects; (2) minor allele frequencies from the 1000 Genomes Project [36], 6500 Exomes Project [37], and the ADNI dataset; and (3) corresponding MutationTaster predictions [38]. We retained all nonsynonymous SNPs, SNPs located in untranslated regions (UTRs), and SNPs with a Regulome DB score less than 4. For each retained SNP, we tested individual associations with case-control status in the 223 controls and 195 cases using the VarStats tool in the Variant Tool Chest (VTC) [39] and subsequently tested their interaction with all SNPs in the other interacting gene using logistic regressions in R.

## 3. Results

### 3.1 Sample and dataset demographics

Sample demographics and minor allele frequencies for rs11136000, rs670139, and rs3865444 are presented for each dataset (Table 1). Eight of the 32 datasets with 3837 cases and 4145 controls passed quality controls for the *CD33-MS4A4E* interaction while seven datasets with 3140 cases and 2713 controls passed for *CLU-MS4A4E.* The remaining datasets were either missing required SNP(s), missing a covariate, or consisted of only controls and could not be included in the analysis. All SNPs passed Hardy-Weinberg equilibrium in all remaining datasets for both cases and controls.

**Table 1.** Sample demographics by dataset. For each dataset the following information 95 is provided: percent cases, females, age, *APOEε4* positive percentage, and minor allele frequencies for rs670139, rs3865444, and rs11136000.

| Study | N | Cases (%) | Females (%) | Age | APOE 4 + (%) | rs670139<br>MAF (T) | rs3865444<br>MAF (A) | rs11136000<br>MAF (T) |
| --- | --- | --- | --- | --- | --- | --- | --- | --- |
| ACT1 | 1858 | 487 (26.2) | 1068 (57.5) | 82.28 | 526 (28.3) | 0.41 | 0.33 | NA |
| ADC2 | 681 | 566 (83.1) | 365 (53.6) | 79.38 | 394 (57.9) | 0.42 | 0.33 | 0.39 |
| ADNI | 371 | 230 (62.0) | 157 (42.3) | 77.82 | 201 (54.2) | 0.45 | 0.31 | 0.37 |
| LOAD | 2965 | 1515 (51.1) | 1882 (63.5) | 78.22 | 1667 (56.2) | 0.43 | 0.31 | 0.38 |
| TARC1 | 388 | 244 (62.9) | 244 (62.9) | 78.96 | 189 (48.7) | 0.43 | 0.32 | 0.41 |
| UMVUMSSM_A | 1058 | 450 (42.5) | 676 (63.9) | 75.48 | 451 (42.6) | 0.43 | 0.31 | 0.38 |
| UMVUMSSM_B | 390 | 135 (34.6) | 236 (60.5) | 73.99 | 118 (30.3) | 0.41 | 0.33 | 0.38 |
| UMVUMSSM_C | 271 | 210 (77.5) | 160 (59.0) | 74.77 | 167 (61.6) | 0.42 | 0.29 | NA |

### 3.2 Homozygous and heterozygous interaction meta-analysis results

The heterozygous interaction between the rs11136000 C/C *(CLU)* and rs670139 G/T *(MS4A4E)* genotypes did not replicate in the independent analysis, though it is suggestive (SF = 1.58, p = 0.07, Figure 1a; Supplemental Table 1). Although the heterozygous interaction did not replicate independently in ADGC, the combined meta-analysis including Cache County is significant (SF = 1.90, p = 0.01, Figure 1a; Supplemental Table 1). The originally reported homozygous *CLU-MS4A4E* interaction between the rs11136000 C/C *(CLU)* and rs670139 G/G *(MS4A4E)* genotypes replicates in the independent meta-analysis (SF = 1.79, p = 0.008, Figure 1b; Supplemental Table 1). The combined meta-analysis is also significant (SF = 2.23, p = 0.0004, Figure 1b; Supplemental Table 1). The individual SNP odds ratios, as previously reported for rs11136000 and rs670139 [21], are 0.83 and 1.09, respectively. The inverted individual SNP odds ratios for rs11136000 and rs670139, are 1.20 and 0.92, respectively. The expected odds ratio for the interaction is 1.20 * 0.92 = 1.10, thus the observed odds ratio is 2.23 * 1.10 = 2.45. Empirical p-values obtained from permutations support the main interaction (ADGC: p = 0.035 with Cache: p = 0.002) and the cPAF for *CLU-MS4A4E* is 8.0. Comparing means to determine whether there is a dosage or dominant effect between the heterozygous and homozygous interactions was not significant (p = 0.22).

**Figure 1.**
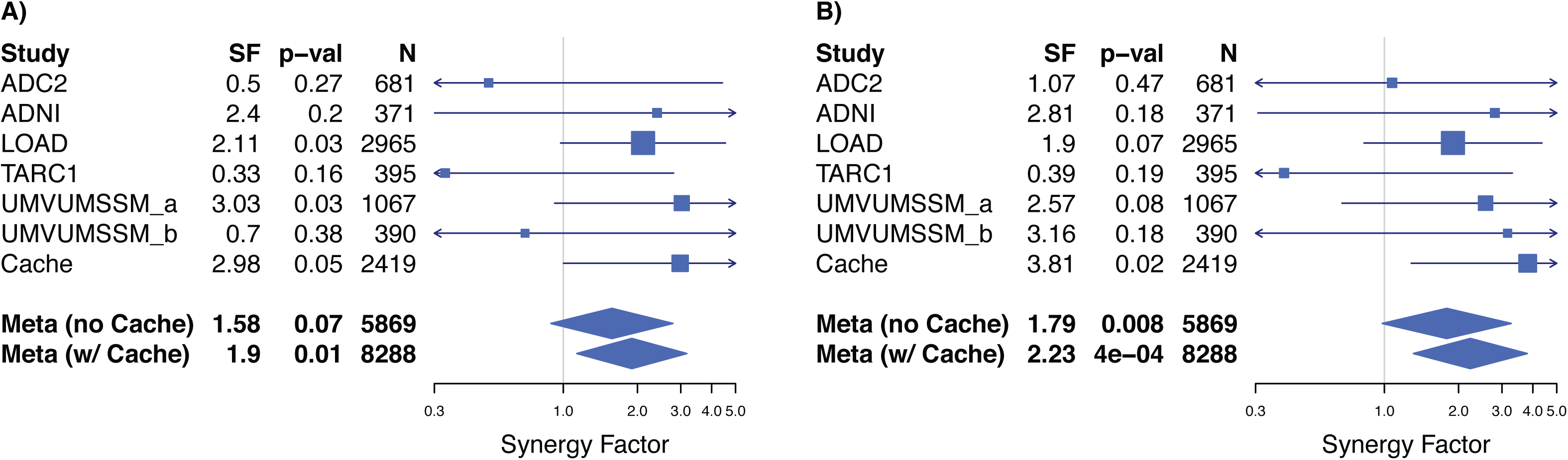
Forest plot showing heterozygous (Panel A) and homozygous (Panel B) *CLU-MS4A4E* interaction replication with potential dominant effect. We tested the original homozygous interaction between the rs11136000 C/C *(CLU;* minor allele = T, MAF = 0.38) and rs670139 G/G *(MS4A4E;* minor allele = T, MAF = 0.38) genotypes, which replicated in ADGC independently (synergy factor = 1.79, p = 0.008, Panel B). We also report the combined metaanalysis including the original Cache County results to present a synergy factor estimate from the largest number of samples possible, which is also significant (synergy factor = 2.23, p = 4e-04, Panel B). We also tested for a dosage or dominant effect based on suggestive evidence in the original Cache County results (Panel A) by testing the heterozygous interaction between the rs11136000 C/C *(CLU)* and rs670139 G/T *(MS4A4E)* genotypes, which did not replicate independently, but is suggestive (synergy factor = 1.58, p = 0.07, Panel A). The combined analysis, including the original Cache County results, is significant (synergy factor = 1.90, p = 0.01, Panel A).

We found an association between the *CLU-MS4A4E* interaction and case-control status in *APOEε4* negative subjects in the combined ADGC and Cache County data (SF = 2.08, p = 0.004, Figure 2a; Supplemental Table 2) that did not exist with *APOEε4* positive subjects (SF = 1.19, p = 0.26, Figure 2b; Supplemental Table 2). The *CD33-MS4A4E* interaction failed to replicate in either the independent or combined meta-analyses (Figures 3a and 3b, Supplemental Table 1).

**Figure 2.**
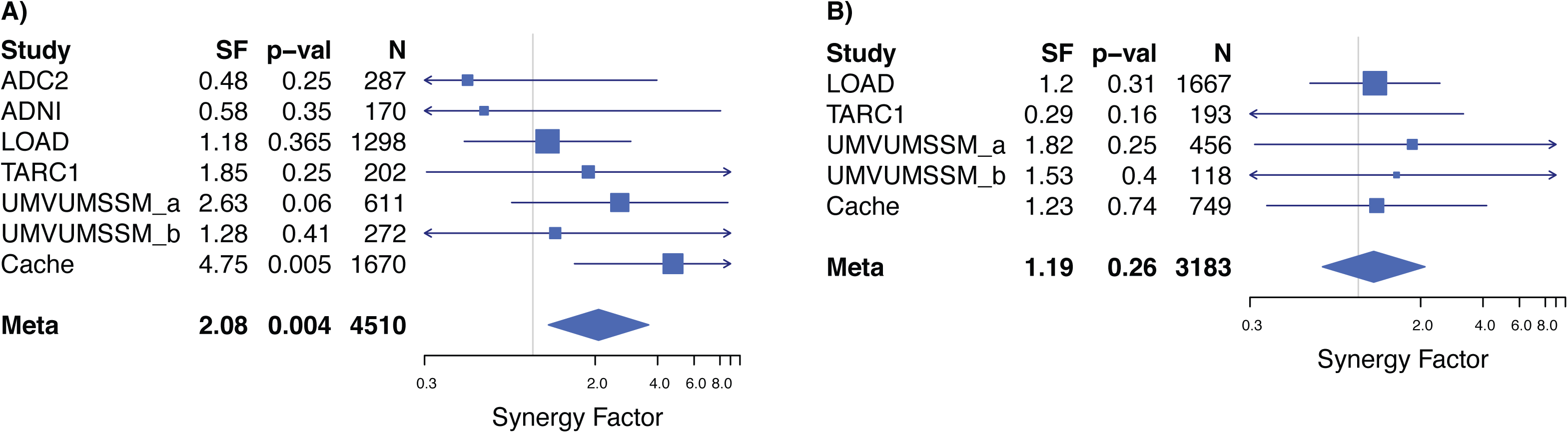
Forest plot showing *APOEε4* negative association with Alzheimer’s disease case-control status. We tested for an *APOEε4* association with the *CLU-MS4A4E* interaction using the Cortina-Borja Synergy Factor Calculator [31]. Specifically, we stratified the combined ADGC and Cache County data by *APOEε4* status and tested for an association between the interaction and case-control status within each stratum. Alleles rs11136000 C and rs670139 G were used as the exposed groups. We found an association in the *APOEε4* negative stratum (synergy factor = 2.08, p = 0.004, Panel A) that did not exist in the *APOEε4* positive stratum (synergy factor = 1.19, p = 0.26, Panel B), suggesting an *APOEε4* effect exists for this interaction.

**Figure 3.**
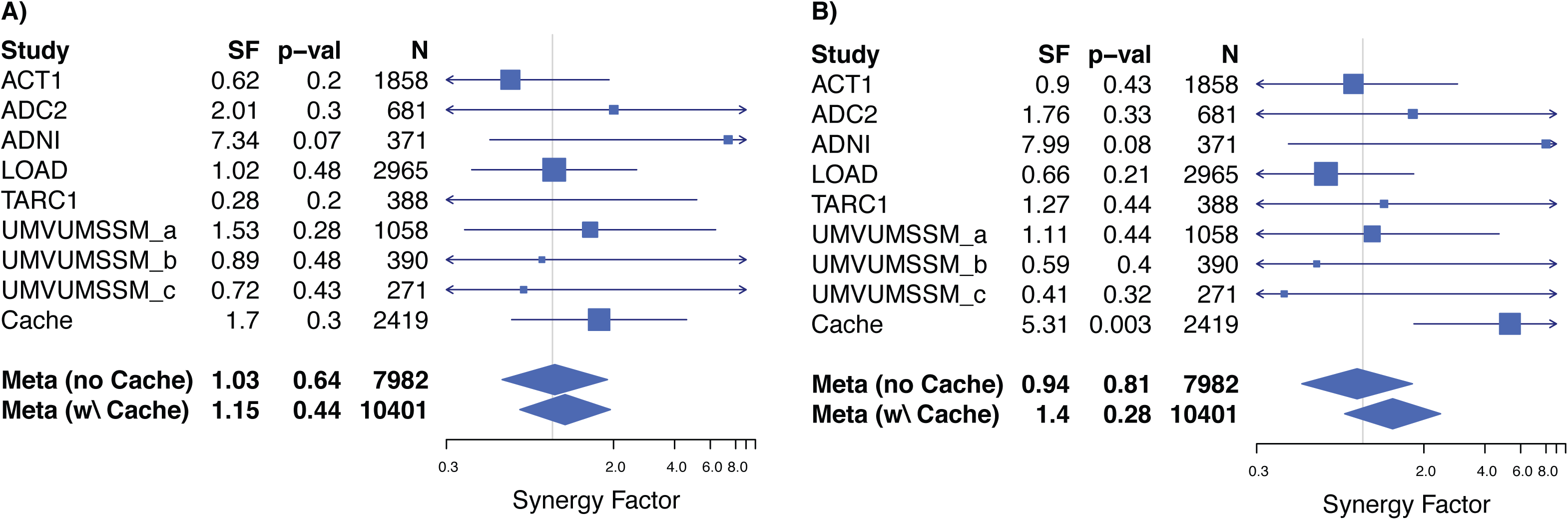
Forest plot showing failed replication for heterozygous (Panel A) and homozygous (Panel B) *CD33-MS4A4E* interaction. We tested the original homozygous interaction between the rs3865444 C/C *(CD33;* minor allele = A, MAF = 0.21) and rs670139 G/G *(MS4A4E;* minor allele = T, MAF = 0.38), which did not replicate in ADGC independently (p = 0.81, Panel B) and was not significant in the combined meta-analysis (p = 0.28, Panel B). We also tested the heterozygous interaction, which also was not significant (without Cache: p = 0.64, Panel A; with Cache: p = 0.44, Panel A).

### 3.3 Exploring causal variants

We explored causal variants in the *CLU* and *MS4A4E* regions using the ADNI whole-genome data. There were 36 and 32 SNPs that fit the inclusion criteria previously described for SNPs near rs11136000 and rs670139, respectively (Supplemental Tables 3 and 4). Most of the SNPs are rare (MAF < 0.01) according to the 1000 Genomes, 6500 Exomes, and ADNI datasets. None of the SNPs were significantly associated with case-control status individually or in the pairwise interactions. We identified two SNPs in *MS4A4E* (rs2081547 and rs11230180) that have a Regulome DB score of ‘ 1f’ and have been shown to modify *MS4A4A* expression [40], the gene upstream from *MS4A4E.* A score of ‘1f’ means they are known to modify expression and are known DNase and transcription factor binding sites.

## 4. Discussion

In this study we attempted to replicate two gene-gene interactions and their association with AD case-control status in the largest dataset used in an epistasis study, to date. The *CD33-MS4A4E* interaction failed to replicate and may have resulted from over-fitting in the Cache County data as previously described by Ebbert et al. [3] Over-fitting happens when a model identifies random data patterns as significant when they are not truly relevant to the question at hand. While there is substantial evidence that CD33 function is related to AD pathways and pathology [16–18], our data do not support an interaction with *MS4A4E* that impacts AD risk.

We replicated the *CLU-MS4A4E* interaction, demonstrated an association in *APOEε4* negative subjects, and reported evidence of a possible dominant effect for *MS4A4E.* The homozygous interaction between rs11136000 *(CLU)* and rs670139 *(MS4A4E)* replicates independently in the ADGC datasets, supporting its validity. To provide synergy factor and odds ratio estimates from the largest number of samples possible, we report the combined meta-analysis synergy factor and odds ratio including the Cache County data. Given the broad sampling and large sample size used for this analysis, our results are likely to be generalizable to other populations of European ancestry. Further investigating this interaction in other ethnic groups is warranted.

Comparing the *CLU-MS4A4E* odds ratio of 2.45 to top AD risk alleles according to AlzGene.org [14] along with *APP, PLD3,* and *TREM2* adds greater perspective. Momentarily ignoring *APOEε4, APP, PLD3,* and *TREM2,* the highest individual odds ratio is from *APOEε2* (OR = 1.61) [3,14] when inverting to its respective risk allele, followed by *ABCA7* (OR = 1.23) [3,14], both of which are dramatically lower than 2.45. Of known AD risk loci, only *APOEε4* (OR = 3.68), *APP* (OR = 5.29), and *TREM2* (OR = 5.05) have ORs greater than 2.45 [3,14,41,42]. The *CLU-MS4A4E* odds ratio is even greater than the *PLD3* Val232Met mutation (OR = 2.10) [43]. These results suggest the *CLU-MS4A4E* interaction may play an important role in AD etiology.

A distinction must be made regarding statistical and biological epistasis, however [22]. While there is evidence that CLU, like CD33, interacts indirectly with MS4A2 [3], little is known about MS4A4E itself and we do not know whether it biologically interacts with CLU. MS4A2 indirectly modifies BCL2L1 activation or expression [3], which physically interacts with CLU. Research suggests CLU prevents amyloid fibrils and other protein aggregation events [44] while MS4A4E may facilitate aggregation as a membrane-spanning protein. Membrane-spanning proteins play diverse roles in cell activity including transport and signaling. Experiments will be required to determine whether there is biological epistasis between CLU and MS4A4E, and whether the interaction affects amyloid fibril formation. Our results indicate further investigative efforts in gene-gene interactions (and protein-protein interactions) may be important to resolve AD etiology.

Comparing means for the effect estimates to assess whether there is a dosage or dominant effect between the *CLU-MS4A4E* heterozygous and homozygous interactions was not significant, suggesting there may be a dominant effect for the rs670139 G allele. A dominant effect has important epidemiologic and heritability implications. Since the *CLU-MS4A4E* interaction increases risk, heterozygous individuals may be at equal risk compared to homozygous individuals.

We found an association between the *CLU-MS4A4E* interaction and case-control status in *APOEε4* negative subjects in the combined ADGC and Cache County data that did not exist with *APOEε4* positive subjects. This potential three-way interaction may provide valuable insight into AD risk and protective factors. A recent paper by Jun et al. [45] found *CLU* has a stronger association in *APOEε4* positive individuals while the region surrounding *MS4A4E* has a stronger association in *APOEε4* negative individuals. Further statistical and biological studies will be necessary to clarify these potential associations. Since all analyses in this study used each gene’s homozygous minor allele as the reference group, the interaction between *CLU-MS4A4E* major alleles is framed as a risk factor, meaning the interaction between the minor alleles is protective. Since the tested heterozygous interaction also increases risk, the protective association may only apply to the interaction between the homozygous minor alleles.

We report several rare potential causal variants linked to rs11136000 or rs670139 with a D’ ≥ 0.99 in the ADNI whole-genome data. No individual variants were significantly associated with AD risk in the ADNI data, but the analysis was likely underpowered with only 240 controls and 202 cases. Two particularly interesting variants, rs11230180 and rs2081547, are known to affect *MS4A4A* expression. We believe further analysis of these variants is necessary to better understand their involvement in AD. Exploring the effects of rs670139, itself, may also be important. Little is known about *MS4A4E,* including the gene’s chromosomal structure. According to gene model XM_011545416.1, rs670139 is in the *MS4A4E* 3’UTR, but other gene models differ. 3’UTR variants can affect transcription and translation.

The cPAF for *CLU-MS4A4E* is 8.0, suggesting there would be an approximate 8% decrease in AD incidence across the population if both major alleles were eliminated. In reality, this estimate is for the causal variants that rs670139 and rs11136000 may be tagging, but the overall effect is nontrivial. Identifying a targeted treatment in the associated pathways could have a significant impact.

A major gap in AD literature to date is the lack of known causal variants. Several SNPs have repeatedly turned up in genome-wide association studies, but the tagSNPs themselves are unlikely to play a direct role in AD etiology. What is more likely is that the tagSNPs are in close linkage disequilibrium with one or more causal variants. We hypothesize two possible explanations: (1) the SNPs are linked to multiple rare variants that drive AD development and progression; or (2) there is another common variant in the region with functional effects that remain unknown. In either case, given the biological complexity of AD and results presented in this study, we believe epistasis plays a critical role in AD etiology. As such, the community must continue to identify and vet these and other interactions that are supported in the literature.

## Acknowledgements

This work was supported by grants from NIH (R01AG11380, R01AG21136, R01AG31272, R01AG042611, R01 AG 042437), the Alzheimer’s Association (MNIRG-11–205368) and the Utah Science, Technology, and Research initiative (USTAR), the Utah State University Agricultural Experiment Station, the McMillian Family Trust, and the Brigham Young University Gerontology Program. The authors thank the participants and staff of the many centers that were involved in data collection for their important contributions to this work. The funders had no role in study design, data collection and analysis, decision to publish, or preparation of the manuscript.The National Institutes of Health, National Institute on Aging (NIH-NIA) also supported this work through the following grants: ADGC, U01 AG032984, RC2 AG036528; NACC, U01 AG016976; NCRAD, U24 AG021886; NIA LOAD, U24 AG026395, U24 AG026390; NIAGADS U24 AG041689; Banner Sun Health Research Institute P30 AG019610; Boston University, P30 AG013846, U01 AG10483, R01 CA129769, R01 MH080295, R01 AG017173, R01 AG025259, R01AG33193; Columbia University, P50 AG008702, R37 AG015473; Duke University, P30 AG028377, AG05128; Emory University, AG025688; Group Health Research Institute, UO1 AG006781, UO1 HG004610, UO1 HG006375; Indiana University, P30 AG10133; Johns Hopkins University, P50 AG005146, R01 AG020688; Massachusetts General Hospital, P50 AG005134; Mayo Clinic, P50 AG016574; Mount Sinai School of Medicine, P50 AG005138, P01 AG002219; New York University, P30 AG08051, M01RR00096, UL1 RR029893, 5R01AG012101, 5R01AG022374, 5R01AG013616, 1RC2AG036502, 1R01AG035137; Northwestern University, P30 AG013854; Oregon Health & Science University, P30 AG008017, R01 AG026916; Rush University, P30 AG010161, R01 AG019085, R01 AG15819, R01 AG17917, R01 AG30146; TGen, R01 NS059873; University of Alabama at Birmingham, P50 AG016582, UL1RR02777; University of Arizona, R01 AG031581; University of California, Davis, P30 AG010129; University of California, Irvine, P50 AG016573, P50, P50 AG016575, P50 AG016576, P50 AG016577; University of California, Los Angeles, P50 AG016570; University of California, San Diego, P50 AG005131; University of California, San Francisco, P50 AG023501, P01 AG019724; University of Kentucky, P30 AG028383, AG05144; University of Michigan, P50 AG008671; University of Pennsylvania, P30 AG010124; University of Pittsburgh, P50 AG005133, AG030653, AG041718, AG07562, AG02365; University of Southern California, P50 AG005142; University of Texas Southwestern, P30 AG012300; University of Miami, R01 AG027944, AG010491, AG027944, AG021547, AG019757; University of Washington, P50 AG005136; University of Wisconsin, P50 AG033514; Vanderbilt University, R01 AG019085; and Washington University, P50 AG005681, P01 AG03991. The Kathleen Price Bryan Brain Bank at Duke University Medical Center is funded by NINDS grant # NS39764, NIMH MH60451 and by Glaxo Smith Kline. Genotyping of the TGEN2 cohort was supported by Kronos Science. The TGen series was also funded by NIA grant AG041232 to AJM and MJH, The Banner Alzheimer’s Foundation, The Johnnie B. Byrd Sr. Alzheimer’s Institute, the Medical Research Council, and the state of Arizona and also includes samples from the following sites: Newcastle Brain Tissue Resource (funding via the Medical Research Council, local NHS trusts and Newcastle University), MRC London Brain Bank for Neurodegenerative Diseases (funding via the Medical Research Council),South West Dementia Brain Bank (funding via numerous sources including the Higher Education Funding Council for England (HEFCE), Alzheimer’s Research Trust (ART), BRACE as well as North Bristol NHS Trust Research and Innovation Department and DeNDRoN), The Netherlands Brain Bank (funding via numerous sources including Stichting MS Research, Brain Net Europe, Hersenstichting Nederland Breinbrekend Werk, International Parkinson Fonds, Internationale Stiching Alzheimer Onderzoek), Institut de Neuropatologia, Servei Anatomia Patologica, Universitat de Barcelona. ADNI Funding for ADNI is through the Northern California Institute for Research and Education by grants from Abbott, AstraZeneca AB, Bayer Schering Pharma AG, Bristol-Myers Squibb, Eisai Global Clinical Development, Elan Corporation, Genentech, GE Healthcare, GlaxoSmithKline, Innogenetics, Johnson and Johnson, Eli Lilly and Co., Medpace, Inc., Merck and Co., Inc., Novartis AG, Pfizer Inc, F. Hoffman-La Roche, Schering-Plough, Synarc, Inc., Alzheimer’s Association, Alzheimer’s Drug Discovery Foundation, the Dana Foundation, and by the National Institute of Biomedical Imaging and Bioengineering and NIA grants U01 AG024904, RC2 AG036535, K01 AG030514. We thank Drs. D. Stephen Snyder and Marilyn Miller from NIA who are ex-officio ADGC members. Support was also from the Alzheimer’s Association (LAF, IIRG-08–89720; MP-V, IIRG-05–14147) and the US Department of Veterans Affairs Administration, Office of Research and Development, Biomedical Laboratory Research Program. P.S.G.-H. is supported by Wellcome Trust, Howard Hughes Medical Institute, and the Canadian Institute of Health Research.

Whole-genome data collection and sharing for this project was funded by the Alzheimer’s Disease Neuroimaging Initiative (ADNI) (National Institutes of Health Grant U01 AG024904) and DOD ADNI (Department of Defense award number W81XWH-12–2–0012). ADNI is funded by the National Institute on Aging, the National Institute of Biomedical Imaging and Bioengineering, and through generous contributions from the following: Alzheimer’s Association; Alzheimer’s Drug Discovery Foundation; Araclon Biotech; BioClinica, Inc.; Biogen Idec Inc.; Bristol-Myers Squibb tooert idCompany; Eisai Inc.; Elan Pharmaceuticals, Inc.; Eli Lilly and Company; EuroImmun; F. Hoffmann-La Roche Ltd and its affiliated company Genentech, Inc.; Fujirebio; GE Healthcare;; IXICO Ltd.; Janssen Alzheimer Immunotherapy Research & Development, LLC.; Johnson & Johnson Pharmaceutical Research & Development LLC.; Medpace, Inc.; Merck & Co., Inc.; Meso Scale Diagnostics, LLC.; NeuroRx Research; Neurotrack Technologies; Novartis Pharmaceuticals Corporation; Pfizer Inc.; Piramal Imaging; Servier; Synarc Inc.; and Takeda Pharmaceutical Company. The Canadian Institutes of Health Research is providing funds to support ADNI clinical sites in Canada. Private sector contributions are facilitated by the Foundation for the National Institutes of Health (www.fnih.org). The grantee organization is the Northern Rev December 5, 2013 California Institute for Research and Education, and the study is coordinated by the Alzheimer’s Disease Cooperative Study at the University of California, San Diego. ADNI data are disseminated by the Laboratory for Neuro Imaging at the University of Southern California.

We further acknowledge Dr. Donald Lehmann and Dr. Mario Cortina-Borja for their kind help with the synergy factor analyses.

This study was made possible by the Texas Alzheimer’s Research and Care Consortium (TARCC) funded by the state of Texas through the Texas Council on Alzheimer’s Disease and Related Disorders. Investigators from TARCC include Baylor College of Medicine: Rachelle Doody MD, PhD, Mimi M. Dang MD, Valory Pavlik PhD, Wen Chan PhD, Paul Massman PhD, Eveleen Darby, Monica Rodriguear RN, Aisha Khaleeq; Texas Tech University Health Sciences Center: Chuang-Kuo Wu MD, PhD, Matthew Lambert PhD, Victoria Perez, Michelle Hernandez; University of North Texas Health Science Center: Thomas Fairchild PhD, Janice Knebl DO, Sid E. O’Bryant PhD, James R. Hall PhD, Leigh Johnson PhD, Robert C. Barber PhD, Douglas Mains, Lisa Alvarez, Rosemary McCallum; University of Texas Southwestern Medical Center: Perrie Adams PhD, Munro Cullum PhD, Roger Rosenberg MD, Benjamin Williams MD, PhD, Mary Quiceno MD, Joan Reisch PhD, Ryan Huebinger PhD, Natalie Martinez, Janet Smith; University of Texas Health Science Center – San Antonio: Donald Royall MD, Raymond Palmer PhD, Marsha Polk; Texas A&M University Health Science Center: Farida Sohrabji PhD, Steve Balsis PhD, Rajesh Miranda, PhD; Essentia Institute of Rural Health: Stephen C. Waring DVM, PhD; University of North Carolina: Kirk C. Wilhelmsen MD, PhD, Jeffrey L. Tilson PhD, Scott Chasse PhD.

## References

[1] James BD, Leurgans SE, Hebert LE, Scherr PA, Yaffe K, Bennett DA. Contribution of Alzheimer disease to mortality in the United States. Neurology 2014;82:1045–50. doi:10.1212/WNL.0000000000000240.

[2] Ridge PG, Mukherjee S, Crane PK, Kauwe JSK, Alzheimer’s Disease Genetics Consortium. Alzheimer’s Disease: Analyzing the Missing Heritability. PLoS ONE 2013;8:e79771. doi:10.1371/journal.pone.0079771.

[3] Ebbert MTW, Ridge PG, Wilson AR, Sharp AR, Bailey M, Norton MC, et al. Population-based Analysis of Alzheimer’s Disease Risk Alleles Implicates Genetic Interactions. Biol Psychiatry 2013. doi:10.1016/j.biopsych.2013.07.008.

[4] Sleegers K, Bettens K, De Roeck A, Cauwenberghe CV, Cuyvers E, Verheijen J, et al. A 22-single nucleotide polymorphism Alzheimer risk score correlates with family history, onset age, and cerebrospinal fluid Aβ42. Alzheimers Dement J Alzheimers Assoc n.d.;0. doi:10.1016/j.jalz.2015.02.013.

[5] Bullock JM, Medway C, Cortina-Borja M, Turton JC, Prince JA, Ibrahim-Verbaas CA, et al. Discovery by the Epistasis Project of an epistatic interaction between the GSTM3 gene and the HHEX/IDE/KIF11 locus in the risk of Alzheimer’s disease. Neurobiol Aging n.d. doi:10.1016/j.neurobiolaging.2012.08.010.

[6] Combarros O, Cortina-Borja M, Smith AD, Lehmann DJ. Epistasis in sporadic Alzheimer’s disease. Neurobiol Aging 2009;30:1333–49. doi:10.1016/j.neurobiolaging.2007.11.027.

[7] Kauwe JSK, Bertelsen S, Mayo K, Cruchaga C, Abraham R, Hollingworth P, et al. Suggestive synergy between genetic variants in TF and HFE as risk factors for Alzheimer’s disease. Am J Med Genet Part B Neuropsychiatr Genet Off Publ Int Soc Psychiatr Genet 2010;153B:955–9. doi:10.1002/ajmg.b.31053.

[8] Naj AC, Jun G, Beecham GW, Wang L-S, Vardarajan BN, Buros J, et al. Common variants at MS4A4/MS4A6E, CD2AP, CD33 and EPHA1 are associated with late-onset Alzheimer’s disease. Nat Genet 2011;43:436–41. doi:10.1038/ng.801.

[9] Harold D, Abraham R, Hollingworth P, Sims R, Gerrish A, Hamshere ML, et al. Genome-wide association study identifies variants at CLU and PICALM associated with Alzheimer’s disease. Nat Genet 2009;41:1088–93. doi:10.1038/ng.440.

[10] Seshadri S, Fitzpatrick AL, Ikram MA, DeStefano AL, Gudnason V, Boada M, et al., CHARGE Consortium, GERAD1 Consortium, EADI1 Consortium. Genome-wide analysis of genetic loci associated with Alzheimer disease. JAMA 2010;303:1832–40. doi:10.1001/jama.2010.574.

[11] Hu X, Pickering E, Liu YC, Hall S, Fournier H, Katz E, et al., the Alzheimer’s Disease Neuroimaging Initiative. Meta-Analysis for Genome-Wide Association Study Identifies Multiple Variants at the BIN1 Locus Associated with Late-Onset Alzheimer’s Disease. PLoS ONE 2011;6:e16616. doi:10.1371/journal.pone.0016616.

[12] Guerreiro RJ, Beck J, Gibbs JR, Santana I, Rossor MN, Schott JM, et al. Genetic Variability in CLU and Its Association with Alzheimer’s Disease. PLoS ONE 2010;5:e9510. doi:10.1371/journal.pone.0009510.

[13] Bettens K, Brouwers N, Van Miegroet H, Gil A, Engelborghs S, De Deyn PP, et al. Follow-up study of susceptibility loci for Alzheimer’s disease and onset age identified by genome-wide association. J Alzheimers Dis JAD 2010;19:1169–75. doi:10.3233/JAD-2010-1310.

[14] Bertram L, McQueen MB, Mullin K, Blacker D, Tanzi RE. Systematic meta-analyses of Alzheimer disease genetic association studies: the AlzGene database. Nat Genet 2007;39:17–23. doi:10.1038/ng1934.

[15] Moebus S, Mecocci P, Del Zompo M, Maier W, Hampel H, Pilotto A et al., with Lambert J-C, Ibrahim-Verbaas CA, Harold D, Naj AC, Sims R, Bellenguez C, et al., European Alzheimer’s Disease Initiative (EADI), Genetic and Environmental Risk in Alzheimer’s Disease (GERAD), Alzheimer’s Disease Genetic Consortium (ADGC), Cohorts for Heart and Aging Research in Genomic Epidemiology (CHARGE). Meta-analysis of 74,046 individuals identifies 11 new susceptibility loci for Alzheimer’s disease. Nat Genet 2013. doi:10.1038/ng.2802.

[16] Morris MC, Evans DA, Johnson K, Sperling RA, Schneider JA, Bennett DA et al., with Bradshaw EM, Chibnik LB, Keenan BT, Ottoboni L, Raj T, Tang A, et al., The Alzheimer Disease Neuroimaging Initiative. CD33 Alzheimer’s disease locus: altered monocyte function and amyloid biology. Nat Neurosci 2013;16:848–50. doi:10.1038/nn.3435.

[17] Griciuc A, Serrano-Pozo A, Parrado AR, Lesinski AN, Asselin CN, Mullin K, et al. Alzheimer’s Disease Risk Gene CD33 Inhibits Microglial Uptake of Amyloid Beta. Neuron 2013;78:631–43. doi:10.1016/j.neuron.2013.04.014.

[18] Karch CM, Jeng AT, Nowotny P, Cady J, Cruchaga C, Goate AM. Expression of Novel Alzheimer’s Disease Risk Genes in Control and Alzheimer’s Disease Brains. PLoS ONE 2012;7:e50976. doi:10.1371/journal.pone.0050976.

[19] Lancaster TM, Brindley LM, Tansey KE, Sims RC, Mantripragada K, Owen MJ, et al. Alzheimer’s disease risk variant in CLU is associated with neural inefficiency in healthy individuals. Alzheimers Dement J Alzheimers Assoc 2014. doi:10.1016/j.jalz.2014.10.012.

[20] Braskie MN, Jahanshad N, Stein JL, Barysheva M, McMahon KL, Zubicaray GI de, et al. Common Alzheimer’s Disease Risk Variant Within the CLU Gene Affects White Matter Microstructure in Young Adults. J Neurosci 2011;31:6764–70. doi:10.1523/JNEUROSCI.5794-10.2011.

[21] Hollingworth P, Harold D, Sims R, Gerrish A, Lambert J-C, Carrasquillo MM et al., Alzheimer’s Disease Neuroimaging Initiative, van Duijn CM, Breteler MMB, Ikram MA, DeStefano AL, Fitzpatrick AL, Lopez O et al., CHARGE consortium, Berr C, Campion D, Epelbaum J, Dartigues J-F, Tzourio C, Alpérovitch A, et al., EADI1 consortium. Common variants at ABCA7, MS4A6A/MS4A4E, EPHA1, CD33 and CD2AP are associated with Alzheimer’s disease. Nat Genet 2011;43:429–35. doi:10.1038/ng.803.

[22] Ebbert M, Ridge P, Kauwe, John. Bridging the gap between statistical and biological epistasis in Alzheimer’s disease. BioMed Res Int In Press.

[23] Kevin L Boehme, Shubhabrata Mukherjee, Paul K Crane, John S K Kauwe. ADGC 1000 Genomes combined data workflow 2014.

[24] Breitner JC, Wyse BW, Anthony JC, Welsh-Bohmer KA, Steffens DC, Norton MC, et al. APOE-epsilon4 count predicts age when prevalence of AD increases, then declines: the Cache County Study. Neurology 1999;53:321–31.

[25] Purcell S, Neale B, Todd-Brown K, Thomas L, Ferreira MAR, Bender D, et al. PLINK: A Tool Set for Whole-Genome Association and Population-Based Linkage Analyses. Am J Hum Genet 2007;81:559–75.

[26] Chang CC, Chow CC, Tellier LC, Vattikuti S, Purcell SM, Lee JJ. Second-generation PLINK: rising to the challenge of larger and richer datasets. GigaScience 2015;4:7. doi:10.1186/s13742-015-0047-8.

[27] Wigginton JE, Cutler DJ, Abecasis GR. A Note on Exact Tests of Hardy-Weinberg Equilibrium. Am J Hum Genet 2005;76:887–93.

[28] Graffelman J, Moreno V. The mid p-value in exact tests for Hardy-Weinberg equilibrium. Stat Appl Genet Mol Biol 2013;12:433–48. doi:10.1515/sagmb-2012-0039.

[29] R Development Core Team. R: A Language and Environment for Statistical Computing. Vienna, Austria: R Foundation for Statistical Computing; 2011.

[30] Willer CJ, Li Y, Abecasis GR. METAL: fast and efficient meta-analysis of genomewide association scans. Bioinformatics 2010;26:2190–1. doi:10.1093/bioinformatics/btq340.

[31] Cortina-Borja M, Smith AD, Combarros O, Lehmann D. The synergy factor: a statistic to measure interactions in complex diseases. BMC Res Notes 2009;2:105. doi:10.1186/1756-0500-2-105.

[32] Lumley T. rmeta: Meta-analysis. 2012.

[33] Boyle AP, Hong EL, Hariharan M, Cheng Y, Schaub MA, Kasowski M, et al. Annotation of functional variation in personal genomes using RegulomeDB. Genome Res 2012;22:1790–7. doi:10.1101/gr,137323.112.

[34] Chang X, Wang K. wANNOVAR: annotating genetic variants for personal genomes via the web. J Med Genet 2012;49:433–6. doi:10.1136/jmedgenet-2012-100918.

[35] Barrett JC, Fry B, Maller J, Daly MJ. Haploview: analysis and visualization of LD and haplotype maps. Bioinformatics 2005;21:263–5. doi:10.1093/bioinformatics/bth457.

[36] Consortium T 1000 GP. An integrated map of genetic variation from 1,092 human genomes. Nature 2012;491:56–65. doi:10.1038/nature11632.

[37] NHLBI GO Exome Sequencing Project (ESP) 2014. http://evs.gs.washington.edu/EVS/.

[38] Schwarz JM, Rödelsperger C, Schuelke M, Seelow D. MutationTaster evaluates disease-causing potential of sequence alterations. Nat Methods 2010;7:575–6. doi:10.1038/nmeth0810-575.

[39] Ebbert MT, Wadsworth ME, Boehme KL, Hoyt KL, Sharp AR, O’Fallon BD, et al. Variant Tool Chest: an improved tool to analyze and manipulate variant call format (VCF) files. BMC Bioinformatics 2014;15:S12. doi:10.1186/1471-2105-15-S7-S12.

[40] Zeller T, Wild P, Szymczak S, Rotival M, Schillert A, Castagne R, et al. Genetics and beyond--the transcriptome of human monocytes and disease susceptibility. PloS One 2010;5:e10693. doi:10.1371/journal.pone.0010693.

[41] Jonsson T, Atwal JK, Steinberg S, Snaedal J, Jonsson PV, Bjornsson S, et al. A mutation in APP protects against Alzheimer/’s disease and age-related cognitive decline. Nature 2012;488:96–9. doi:10.1038/nature11283.

[42] Guerreiro R, Wojtas A, Bras J, Carrasquillo M, Rogaeva E, Majounie E, et al. TREM2 Variants in Alzheimer’s Disease. N Engl J Med 2013;368:117–27. doi:10.1056/NEJMoal211851.

[43] Sassi C, Bras J, Gibbs JR, Hernandez DG, Lupton MK, Powell J et al., with Cruchaga C, Karch CM, Jin SC, Benitez BA, Cai Y, Guerreiro R, et al., UK Brain Expression Consortium (ukbec). Rare coding variants in the phospholipase D3 gene confer risk for Alzheimer/’s disease. Nature 2014;505:550–4. doi:10.1038/nature12825.

[44] Yerbury JJ, Poon S, Meehan S, Thompson B, Kumita JR, Dobson CM, et al. The extracellular chaperone clusterin influences amyloid formation and toxicity by interacting with prefibrillar structures. FASEB J Off Publ Fed Am Soc Exp Biol 2007;21:2312–22. doi:10.1096/fj.06-7986com.

[45] Jun G, Ibrahim-Verbaas CA, Vronskaya M, Lambert J-C, Chung J, Naj AC, et al. A novel Alzheimer disease locus located near the gene encoding tau protein. Mol Psychiatry 2015. doi:10.1038/mp.2015.23.

